# Phosphate Limitation Modulates *Vibrio cholerae* Outer Membrane Vesicle Formation, Composition and Toxicity

**DOI:** 10.1101/2025.11.13.688205

**Authors:** Matheus Luchetta da Fonseca, Livia Carvalho Barbosa, Beatriz Ferreira de Carvalho Patricio, Wellington da Silva Ferreira, Carolina Neumann Keim, Cedric Delporte, Pierre van Antwerpen, Jean-Marie Ruysschaert, Paulo Mascarello Bisch, Wanda Maria Almeida von Kruger

**Affiliations:** Laboratório de Física-Biológica, Instituto de Biofísica Carlos Chagas Filho, Universidade Federal do Rio de Janeiro, Rio de Janeiro 21941-902, Brasil; Laboratório de Inovação e Tecnologia Farmacêutica, Departamento de Ciências Fisiológicas −Farmacologia, Instituto Ciências Biomédicas, Universidade Federal do Estado do Rio de Janeiro, Rio de Janeiro 20211-040, Brasil; Centro Nacional de Bioimagem, Universidade Federal do Rio de Janeiro, Rio de Janeiro 21941-902, Brasil; Instituto de Microbiologia Paulo de Góes, Universidade Federal do Rio de Janeiro - UFRJ, Av. Carlos Chagas Filho, 373, Cidade Universitária, 21941-902, Rio de Janeiro, RJ, Brazil; RD3-Pharmacognosy, Bioanalysis and Drug Discovery and Analytical Platform, Faculty of Pharmacy, Université Libre de Bruxelles, 1050 Brussels, Belgium; Structure and Function of Biological Membranes Laboratory, Université Libre de Bruxelles, 1050 Brussels, Belgium

**Keywords:** *Vibrio cholerae*, outer membrane vesicles (OMVs), phosphate limitation, PhoB/PhoR two-component system, virulence

## Abstract

*Vibrio cholerae* inhabits phosphorus-poor aquatic environments and host intestine, where it expresses genes regulated by the PhoB/PhoR two-component system in response to inorganic phosphate (Pi) limitation. Like other Gram-negative bacteria, *V. cholerae* releases outer membrane vesicles (OMVs), which carry proteins, lipids, and nucleic acids that contribute to adaptation, survival, and pathogenesis. Here, we investigated how Pi availability affects OMV production, composition, and toxicity in the pandemic strain N16961 and its Δ*phoB* mutant. Using transmission electron microscopy, atomic force microscopy, and nanoparticle tracking analysis, we show that OMV size remains constant (∼140 nm) across conditions, but production is significantly increased under Pi limitation in a PhoB-dependent manner. Proteomic and lipidomic analyses revealed selective packaging of PhoB-regulated proteins involved in phosphate metabolism, stress response, carbon metabolism, and toxicity, as well as enrichment in phosphorus-free ornithine lipids under low Pi. Functional assays in *Galleria mellonella* demonstrated that OMVs from N16961 under Pi limitation are highly toxic, whereas OMVs from high-Pi or Δ*phoB* cultures exhibit minimal lethality. Our findings indicate that phosphate limitation acts as an environmental cue that shapes OMV composition, enhancing both bacterial survival and pathogenic potential. This study highlights OMVs as dynamic vehicles integrating adaptation to low Pi concentration, stress adaptation, and toxicity in *V. cholerae*.

## Introduction

Cholera is a devastating diarrheal disease caused by *Vibrio cholerae* [1], and its transmission occurs mainly through consumption of contaminated food and/or water. Epidemiological studies estimate that about 1.3 billion people live in endemic countries and are at risk of contracting the disease, a number that may increase due to climate change and environmental catastrophes [2]. Annually, according to WHO, it is estimated that 1.3 to 4 million cases of cholera occur in 69 endemic countries, leading to 143,000 deaths from the disease [3]. Bacteria of the genus *Vibrio* are sensitive to acids, so most ingested cells die in the stomach. Those that manage to survive reach the small intestine, where, after crossing the mucosal layer and reaching the intestinal epithelium, *V. cholerae* cells exhibit reduced motility. This reduction facilitates colonization of the epithelium, biofilm formation, and the production of key virulence factors involved in the infection process, such as the TCP pilus and cholera toxin (CT) [4].

Bacteria’s ability to survive and grow in different environments depends on gene expression regulation. Aquatic and terrestrial environments inhabited by *V. cholerae* are usually deficient in phosphorus [5], an essential component of living beings’ essential components such as nucleic acids, phospholipids, and lipopolysaccharides. The main source of phosphorus by bacteria is inorganic phosphate (Pi) [6], and bacterial species survival, including *V. cholerae*, is directly dependent on its availability [7]. In several bacterial species, including *V. cholerae*, growth in Pi-deficient media leads to activation of the PhoB/PhoR two-component system to increase Pi uptake and enable bacterial survival. The PhoR protein is transmembrane and has a signal sensor domain, while PhoB is a cytoplasmic transcriptional regulator, which affects the signal response [8].

Bacteria use different mechanisms to overcome stressful situations, adapt, and survive in hostile environments. One of these mechanisms is the release of outer membrane vesicles (OMVs) - which are produced by a wide variety of Gram-negative bacteria, including *Vibrio* species [9], and are constantly released from cell surface during bacterial growth [10]. The increase in its release can be stimulated by environmental stress, such as temperature, nutrient deficiency, among others [11]. OMVs are spherical, ranging from 10 to 300 nm in diameter, and are composed mainly of phospholipids, proteins, and lipopolysaccharides (LPS) from the outer membrane, but may also contain periplasm and cytoplasm proteins, in addition to DNA and RNA [10].

Distinct mechanisms have been described in recent decades for OMVs formation by Gram-negative bacteria. The most recent hypothesis for this process is phospholipids (PL) accumulation on the outer membrane outer surface, where only LPS molecules are normally found [12]. This causes bulging in the outer face region of the membrane and OMVs secretion [12,13]. This process has been demonstrated in *Haemophilus influenzae*, *V. cholerae*, and *E. coli*, indicating that this is a conserved mechanism among several species [11]. Incorporation of molecules into OMVs is an important strategy because it confers benefits to the packaged biomolecules, such as protection against degradation by other bacteria, host and environmental factors. It can also maintain a favorable microenvironment for enzymatic activities and greater potential for long-distance delivery [14].

OMVs are important in both physiology and bacterial pathogenesis, acting in the interaction between bacteria, bacteria-host, bacteria-environment, and in response to multiple stresses for adaptation and survival, among others [15]. Some OMVs components exert specialized functions under certain environmental conditions, such as quorum sensing (QS) communication, antibiotic resistance, defense against bacteriophages and members of microbial communities, nucleic acid transfer, horizontal gene transfer, toxin release, factors of virulence and immunomodulators [14,16,17].

In this work, we investigate the composition of outer membrane vesicles released by *Vibrio cholerae* N16961 strain serogroup O1 and its mutant Δ*phoB* grown under Pi limitation or not, to assess the contribution of the PhoB/PhoR regulatory system to this process. Our objective is to determine how changes in OMV release and cargo are influenced by phosphate availability, and how the components of the vesicles may contribute to bacterial adaptation, survival, and pathogenicity.

## Results and Discussion

### Characterization of OMVs released by *V. cholerae* N16961 and Δ*phoB* under abundance and phosphate limitation

The OMVs released by N16961 and Δ*phoB* strains grown in high and low phosphate were analyzed by transmission electron microscopy (TEM) (Fig. 1A-C). They consisted of spherical vesicles of variable sizes, forming chains and aggregates (Fig. 1B-C). Interestingly, interconnected outer membrane vesicles chains have been reported for many bacterial species, including *Vibrio* species, and it has been shown that, in many cases, they function as bridges in cell-cell interactions to transfer membrane proteins and other molecules between cells [18]. The low amounts of debris and the absence of intact cells confirmed the high purity of the OMVs preparations (Figure 1B and C).

**Fig 1.**
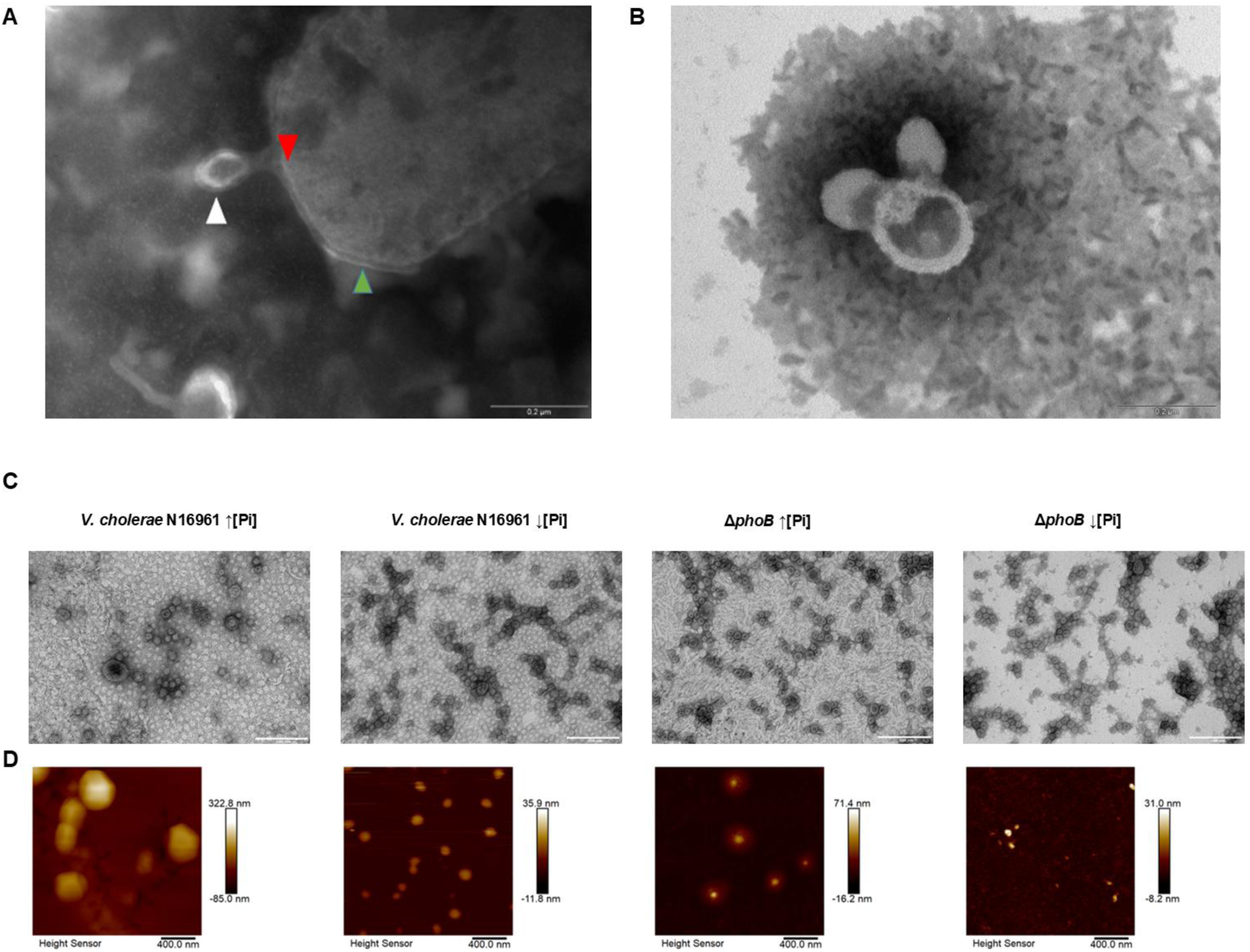
Morphological characterization of OMVs from *Vibrio cholerae* N16961 and ΔphoB under phosphate limitation. A) Representative TEM image of an OMV budding from the outer membrane in *V. cholerae* strain N16961 grown in low Pi - red arrow showing the inner membrane, green arrow the outer membrane and the white arrow a budging vesicle. B) TEM image of vesicles from a purified preparation of OMVs from strain N16961 under low Pi. C) TEM images from purified OMVs. D) AFM images with 2 µm size obtained on a Bruker Icon microscope from purified OMVs.

TEM analysis also permitted us to observe a budding vesicle on *V. cholerae* strain N16961 cell surface (Fig. 1A), showing that OMV formation starts by outer membrane local curvature and budding, followed by its release, a mechanism that seems to be highly conserved among Gram-negative species [12,19].

Both TEM and AFM images revealed agglomerate structures composed of individual OMVs with clearly defined boundaries between vesicles (Fig. 1C-D). AFM imaging showed OMV sizes consistent with TEM observations and further revealed the presence of a distinct extracellular layer on the bacterial surface under high phosphate (Pi) conditions (Fig. D). Specifically, N16961 exhibited a layer of ∼37 ± 6 nm, whereas the Δ*phoB* displayed a thicker layer of ∼113 ± 37 nm. This extracellular layer was absent when bacteria were grown under low Pi (Fig. D).

### OMVs released by *V. cholerae* N16961 and Δ*phoB* grown under Pi limitation and abundance have similar size distribution ranges but are produced in different amounts

The effects of the *phoB* gene mutation on the size and production rate of OMVs by the pandemic wild-type *V. cholera* strain N16961, under inorganic phosphate abundance and limitation (Pi) were analyzed.

Nanoparticle Tracking Analysis (NTA) showed that strains N16961 and Δ*phoB*, released homogeneous populations of spherical OMVs with an average diameter of 140.5 nm under both conditions (Fig. 2A).

**Fig 2.**
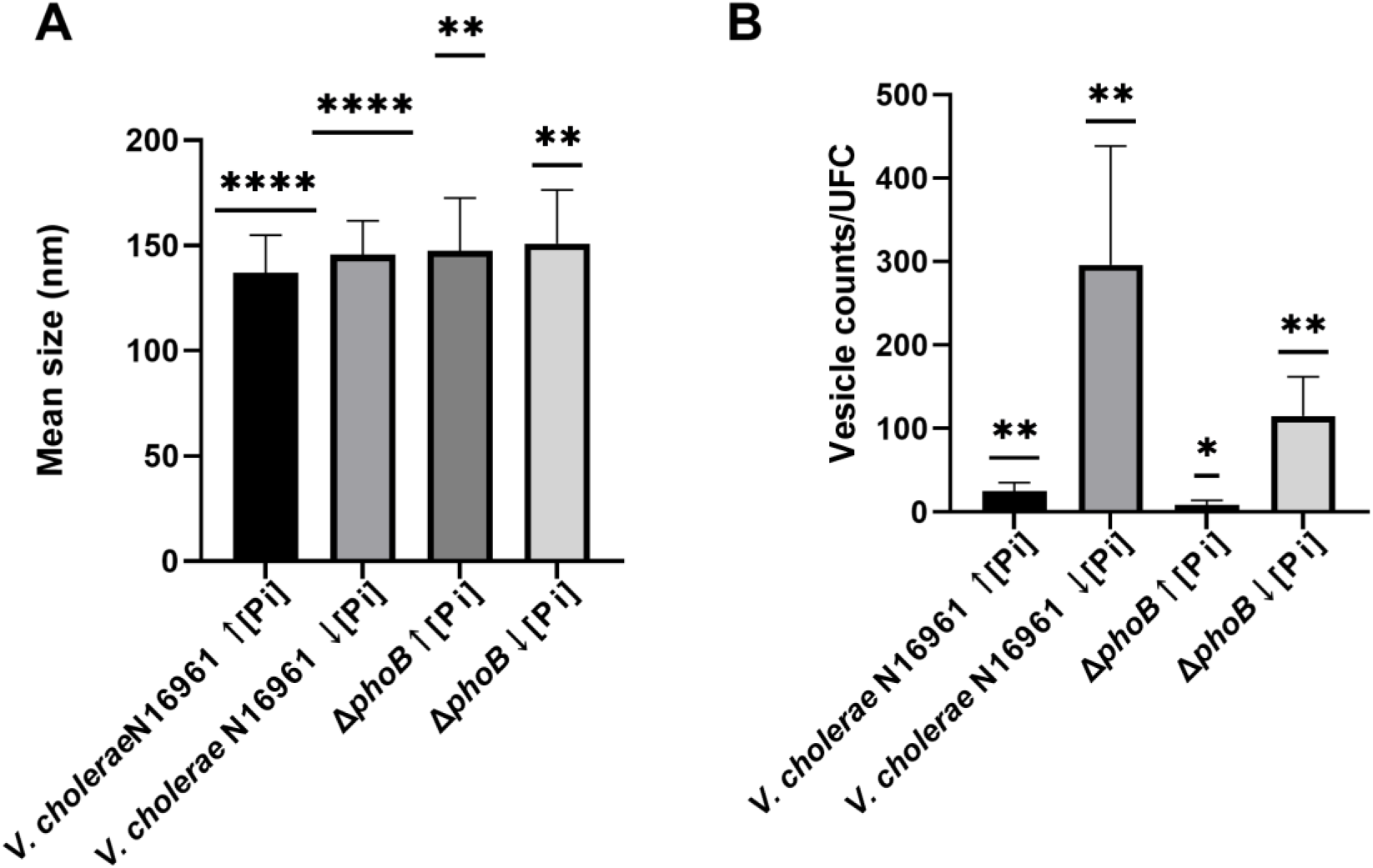
Nanoparticle Tracking Analysis (NTA) of OMVs produced by *V. cholerae* N16961 and ΔphoB under high and low phosphate. Mean diameters of purified OMVs from five independent cultures of *V. cholerae* N16961 and ΔphoB grown under high and low Pi. Both strains produce vesicles of similar size distributions, but in significantly different quantities under Pi limitation.

These results showed that mutation in the *phoB* gene and/or growth under Pi limitation (activation of the PhoB/R system) did not cause significant changes in the size of OMVs produced from *V. cholera* N16961 (Fig. 2A). However, the amount of OMVs released by the strains varied with the Pi concentration on the culture medium and was affected by the mutation in the *phoB* gene (Fig. 2B). Under Pi abundance, strains N16961 and Δ*phoB* produced similar and very low amounts of OMVs/CFU (Fig. 2B). On the other hand, Pi limitation caused a significant increase in OMVs/UFC production by both strains (Fig. 2B). Under this condition, Δ*phoB* released about 100 OMVs/UFC, whereas the wild type N16961 produced 300 OMVs/UFC (Fig. 2B).

The increased OMV production by the wild-type strain N16961 under Pi deficiency compared to Pi abundance is consistent with the higher vesiculation observed in many Gram-negative bacterial species under stress [13,21,22]. To ensure that this difference was not simply due to variations in bacterial cell numbers, OMVs were isolated from cultures normalized by total cell counts.

### OMVs released by *V. cholerae* N16961 under Pi limitation are toxic to the *Galleria mellonella* larvae model in a PhoB/PhoR-dependent manner

Under Pi limitation, the *V. cholera* PhoB/PhoR system activates the Pho regulon, which includes genes involved in various cellular processes, including pathogenesis.

To investigate whether OMVs released by the *V. cholera* strain N16961 and its *phoB* mutant in Pi abundance or limitation carried virulence factors, purified OMVs were tested using larvae of the greater wax moth *Galleria mellonella* as the toxicity model [24,25]. Therefore, *G. mellonella* larvae were inoculated with purified OMVs released by the wild-type strain N16961 and Δ*phoB* in media with high and low Pi. After inoculation, the larvae were kept at 37 ^◦^C for 72 h, under protection from light, and mortality was evaluated every 12 h.

The results showed no death among larvae in the two control groups (Untouched larvae and inoculated with PBS) in the experiment’s timeline. However, among the inoculated groups, larval death was time-dependent and started at 24 h post inoculation (p.i.). The survival outcome of inoculated larvae varied depending on the OMVs origin. Those released by Δ*phoB* in both conditions induced a negligible larval mortality rate (∼ 5% in 72 h; p < 0.05) (Fig. 3).

**Fig 3.**
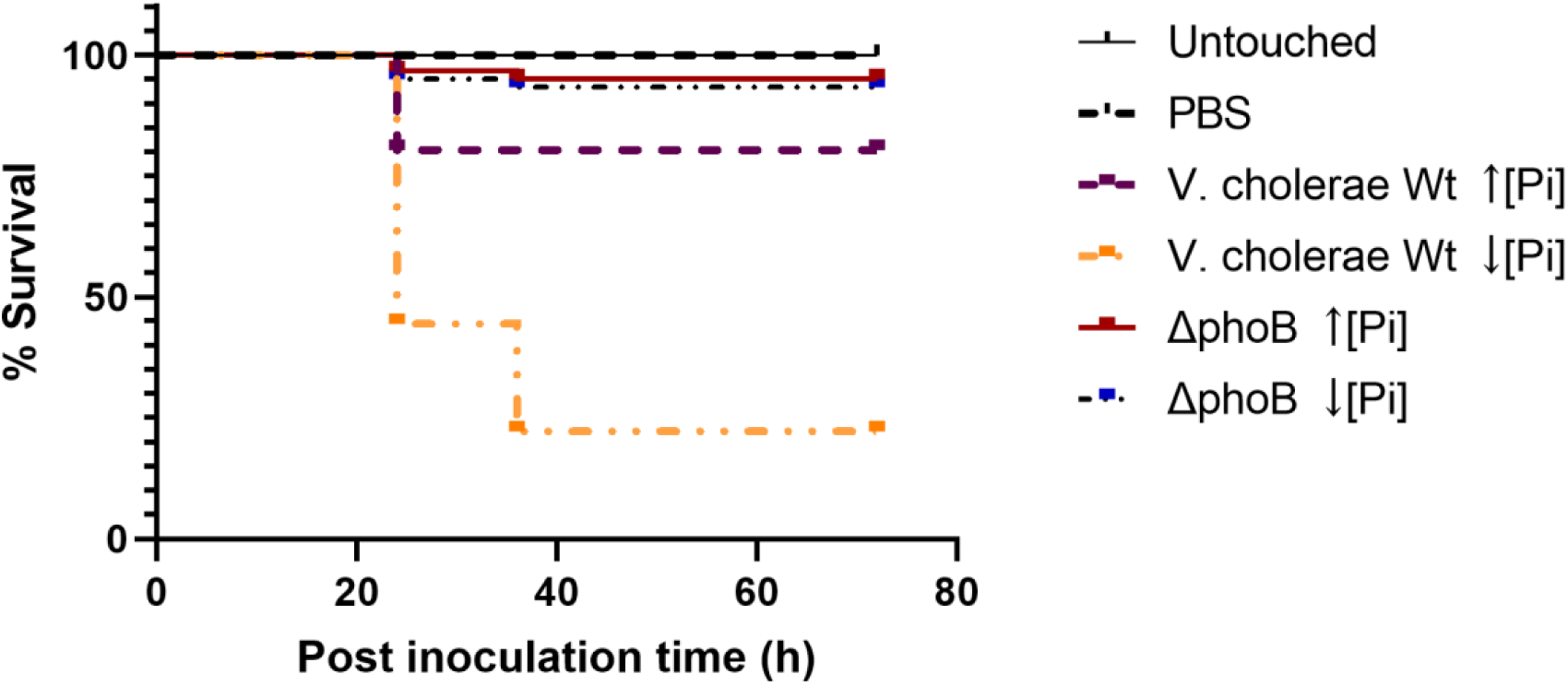
OMVs produced under low phosphate increase lethality in *Galleria mellonella.* Survival rate of *G. mellonella* larvae inoculated with OMV suspensions released by *V. cholerae* N16961 and ΔphoB grown under high or low Pi. Controls: larvae uninoculated or injected with 10 μL of sterile PBS. Purified OMVs (n>10 per experiment, total n=60). Data analyzed by Kaplan–Meier test (*p* < 0.05).

In contrast, OMVs released by the wild-type strain N16961 in high Pi significantly increased larval mortality (20%) 24 h p.i., without further change. However, the highest larval mortality (80 %) occurred among those inoculated with N16961 OMVs from low Pi cultures. After 24 h p.i. about 50% of the inoculated larvae were dead, and 12h later, the mortality reached 80 %, without further change up to 72 h p.i. (p<0.05) (Fig. 3).

These results showed that OMVs released by *V. cholerae* N16961 under Pi limiting conditions can specifically carry toxic compounds, unlike OMVs obtained under the other tested conditions. This finding suggests that phosphate availability may act as a regulatory signal influencing the toxicity of OMVs. While previous studies have established a link between the Pho regulon and *V. cholerae* virulence, our data expand this knowledge by showing that Pi limitation not only regulates gene expression but also shapes the vesicular cargo, potentially contributing to host-pathogen interactions and virulence.

### Proteomic analysis of the OMVs liberated by N16961 and Δ*phoB* under limitation and abundance of inorganic phosphate showed individual, shared, and core proteins

OMV proteins released by N16961 and Δ*phoB*, in high and low Pi, were analyzed by liquid chromatography coupled to high resolution tandem mass spectrometry.. In total, 370 unique proteins were identified over the four conditions (Supplementary material X). We found 125 proteins in OMVs released by N16961 in high Pi and 95 in those from low Pi. For the mutant strain, 189 proteins were identified in the OMVs released in high Pi and 277 in those generated in low Pi. The proteins from each OMV type were separated according to functional categories (COGs). The total number of proteins in each OMV type was defined as 100%, and the percentage of proteins in each COG was determined (Fig. 4).

**Fig 4.**
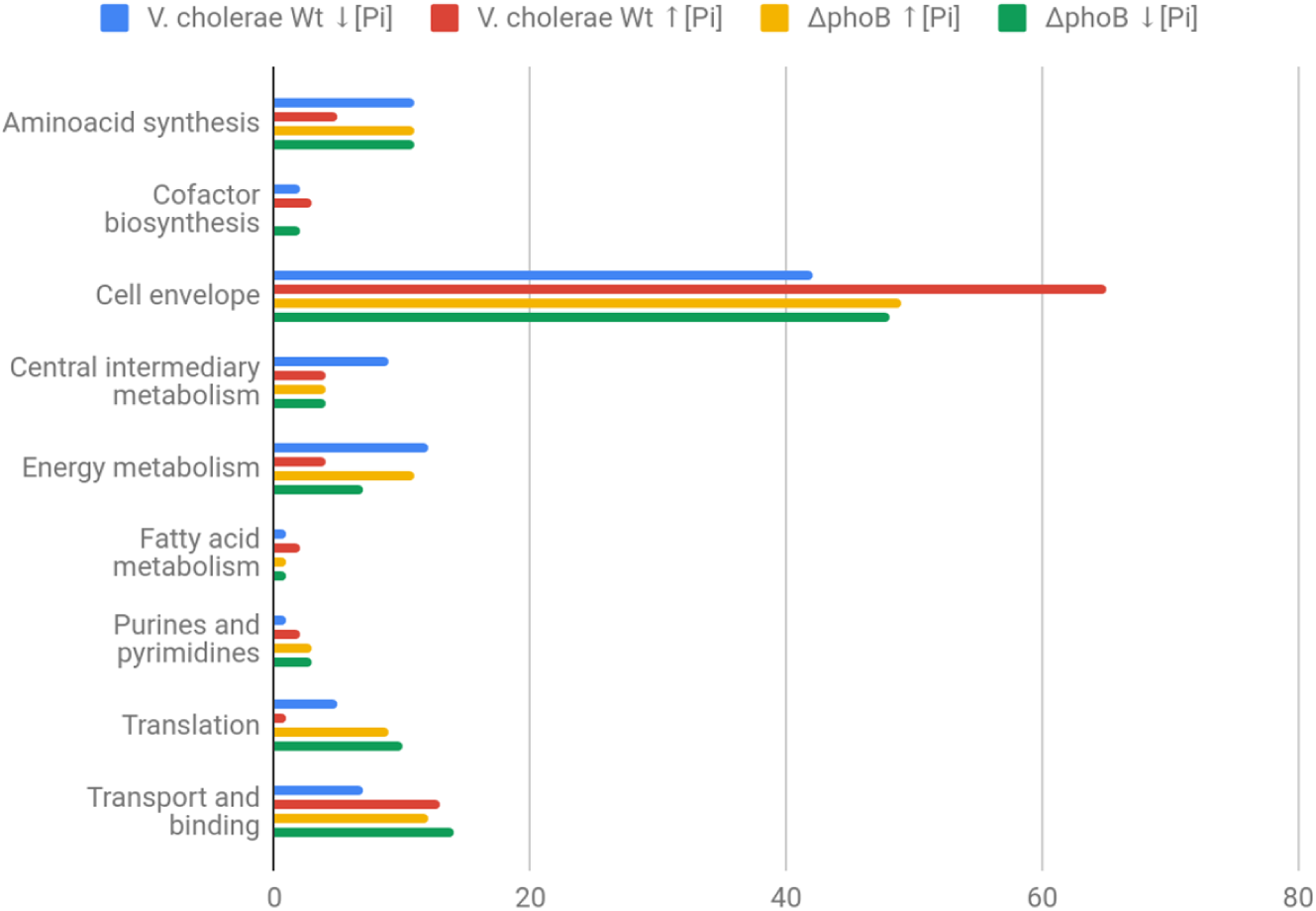
Functional categorization of proteins identified in OMVs of *V. cholerae* under phosphate limitation and abundance. Proteins identified by LC–MS/MS were analyzed using Peaks Studio X against the UniProt database for *V. cholerae* and classified into functional categories according to Clusters of Orthologous Groups (COGs). Percentages represent the proportion of proteins in each COG category relative to the total identified under each condition.

The sum of all identified proteins was submitted to Venn analysis, which identified shared and exclusive proteins of these vesicles (Fig. 5A). Venn analysis revealed 50 proteins that were shared between OMVs derived from strain N16961 and those from the Δ*phoB*. As for unique proteins, the OMVs from N16961 generated in high Pi have 12 (∼10% of those identified) and those from low Pi, 24 (∼25% of those identified). In the case of Δ*phoB*, the OMVs released in high Pi have 43 (∼23% of those identified) and those from low Pi, 114 (∼41%).

**Fig 5.**
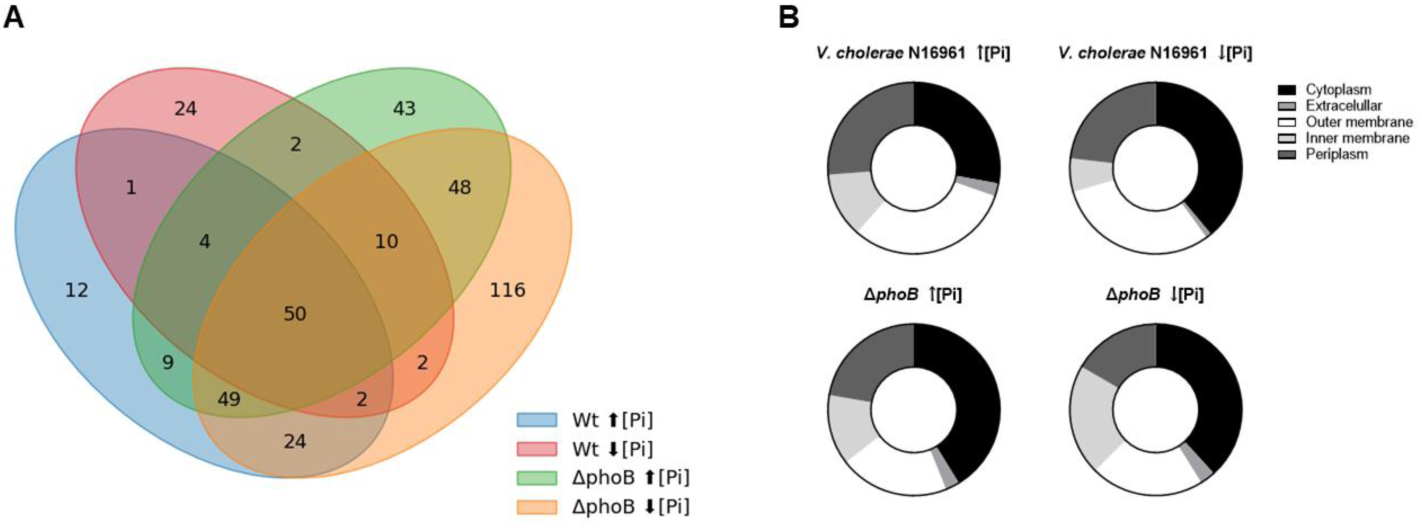
Proteomic comparison of OMVs from *V. cholerae* N16961 and ΔphoB under different phosphate conditions. (A) Venn diagram showing the number of unique and shared proteins in OMVs from V. cholerae N16961 and ΔphoB grown under high and low Pi conditions. (B) Subcellular localization of proteins identified in OMVs from N16961 and ΔphoB under high and low Pi. Cellular localization was predicted using the CELLO program (http://cello.life.nctu.edu.tw/).

Except for OMVs released by the wild-type strain N16961 in high Pi, enriched in similar amounts of outer membrane, periplasm, and cytoplasmic proteins, the other three OMV groups had proteins from the cytoplasm as their main components. In addition, the four OMV types carried small amounts of both internal membrane and extracellular proteins (Fig. 5B). The presence of cell membrane-associated, cytoplasmic, and periplasmic proteins in the four OMVs types supports the observations that OMV components derive from multiple bacterial compartments [26], but it depends on the bacterial strains analyzed and growth conditions.

Therefore, the distinct cargo proteins in the four OMV types can be related to the genetic difference between the *V. cholerae* wild-type N16961 and its Δ*phoB* strain and also to the growth conditions under high and low Pi, as previously reported in the proteomic study of the bacteria, which showed that the Pi limitation leads to differential expression of PhoB dependent/independent proteins, influencing essential factors for *V. cholerae* adaptation and virulence [27]. In particular, the wild-type strain grown under Pi limitation induced Pho regulon proteins involved in phosphate metabolism like PstS and alkaline phosphatase, while the Δ*phoB* relied mainly on stress related proteins, such as heat-shock proteins, outer membrane components and detoxification enzymes. These findings highlight that Pi availability not only modulates protein composition at cellular level but may also influence the specific composition of secreted vesicular cargo, with potential benefits for bacterial survival and pathogenicity.

### OMVs released by *V. cholera* N16961 under Pi limitation are vehicles for specific proteins involved in metabolism, transport, stress response, and gene expression regulation - some of these are involved in bacterial virulence

OMVs are produced by all Gram-negative bacteria and function as delivery systems for soluble molecules and insoluble compounds, which they release into the environment, contributing to bacterial survival and virulence. Among the proteins identified in OMVs N16961 strain under Pi limitation, some are products of genes regulated by the PhoB/PhoR system (VON KRUGER et al., 2006). We analyzed the 24 proteins (Table 1) that were identified exclusively in OMVs released by the N16961 strain under Pi limitation, in an attempt to elucidate their possible roles in bacterial survival under stress conditions, nutrient acquisition, and the lethal effect of the vesicles on Galleria mellonella larvae.

**Table 1.**
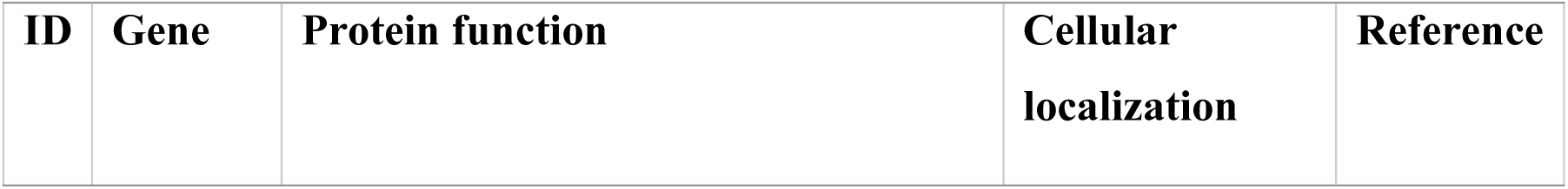
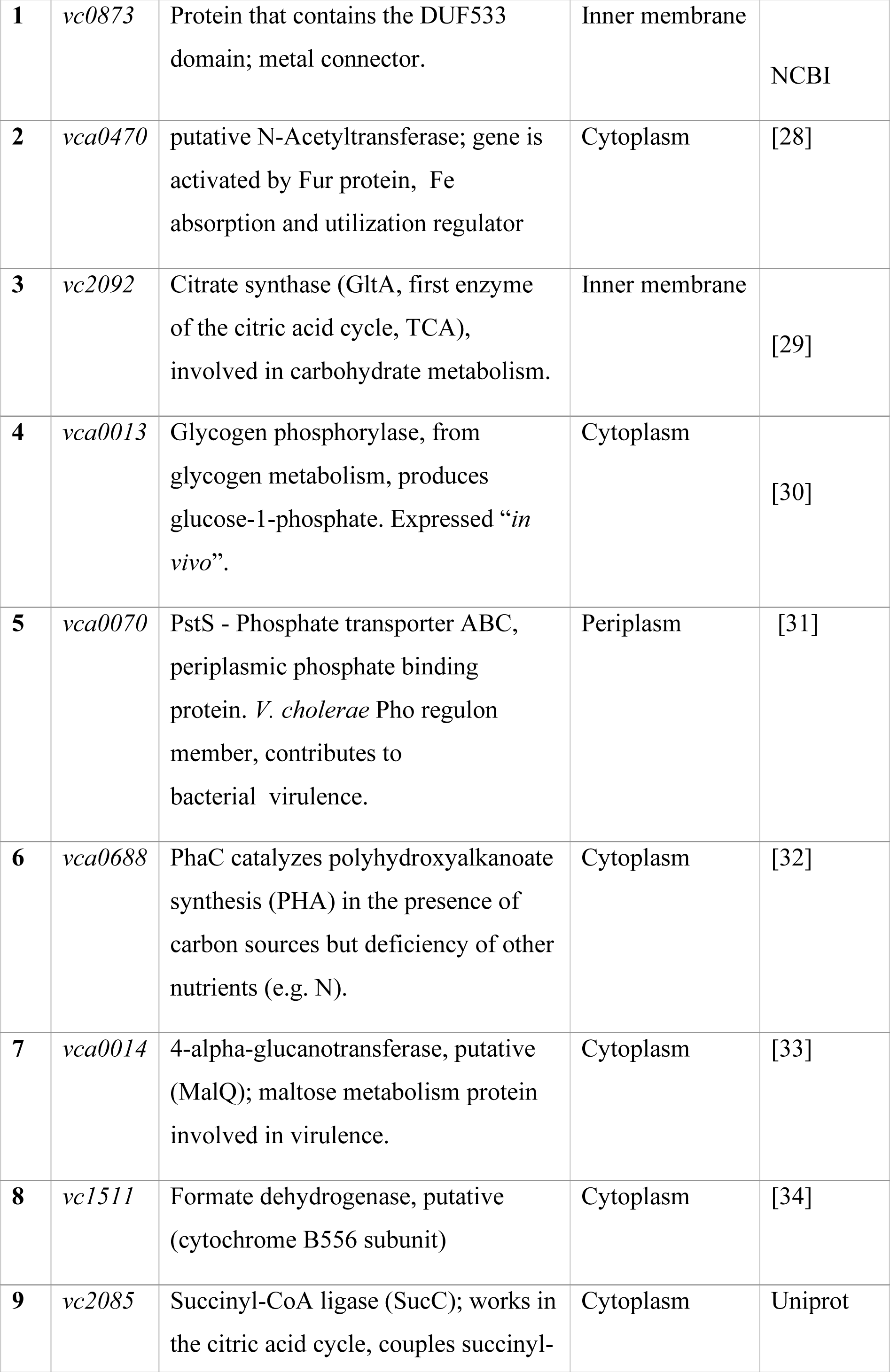
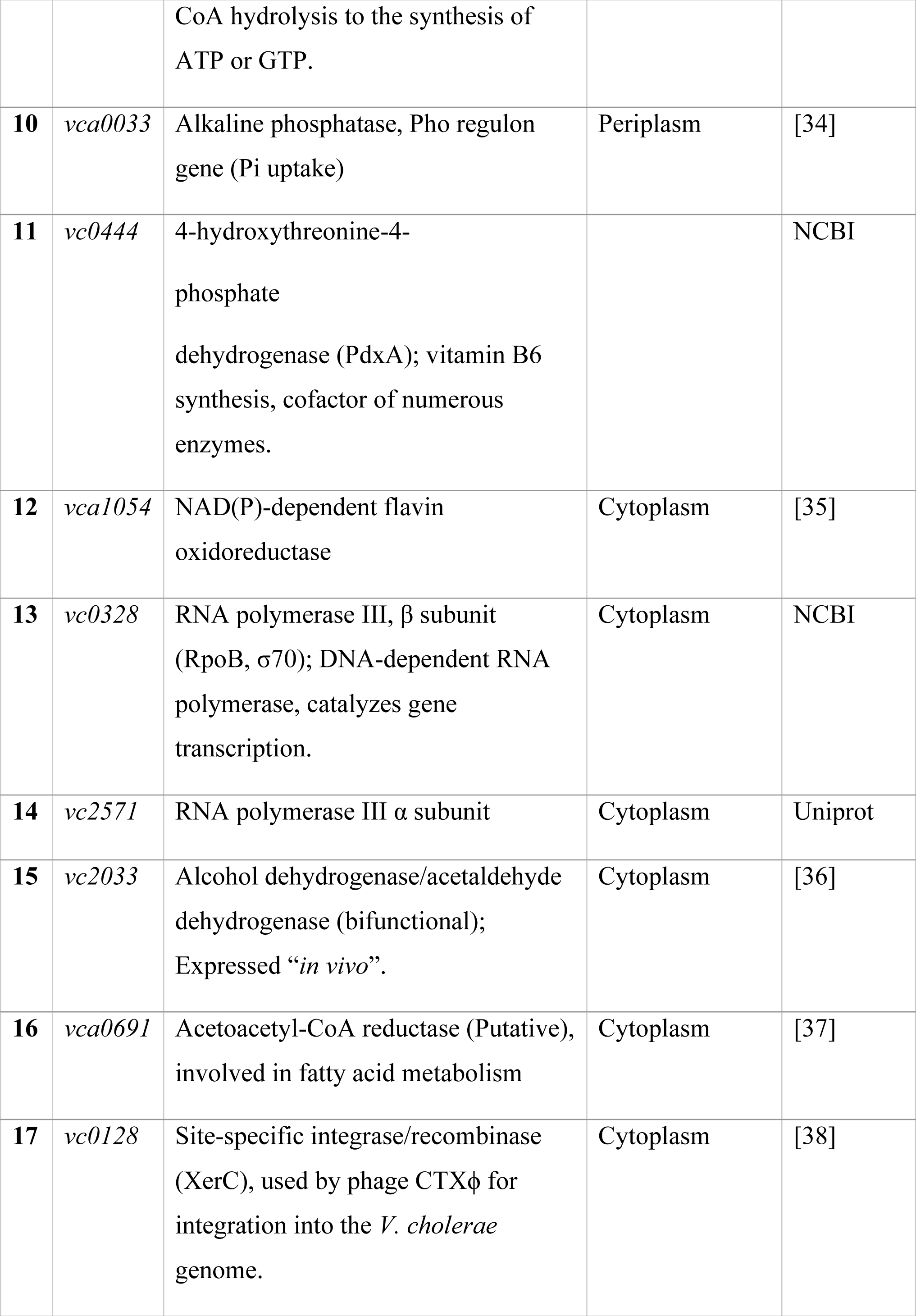
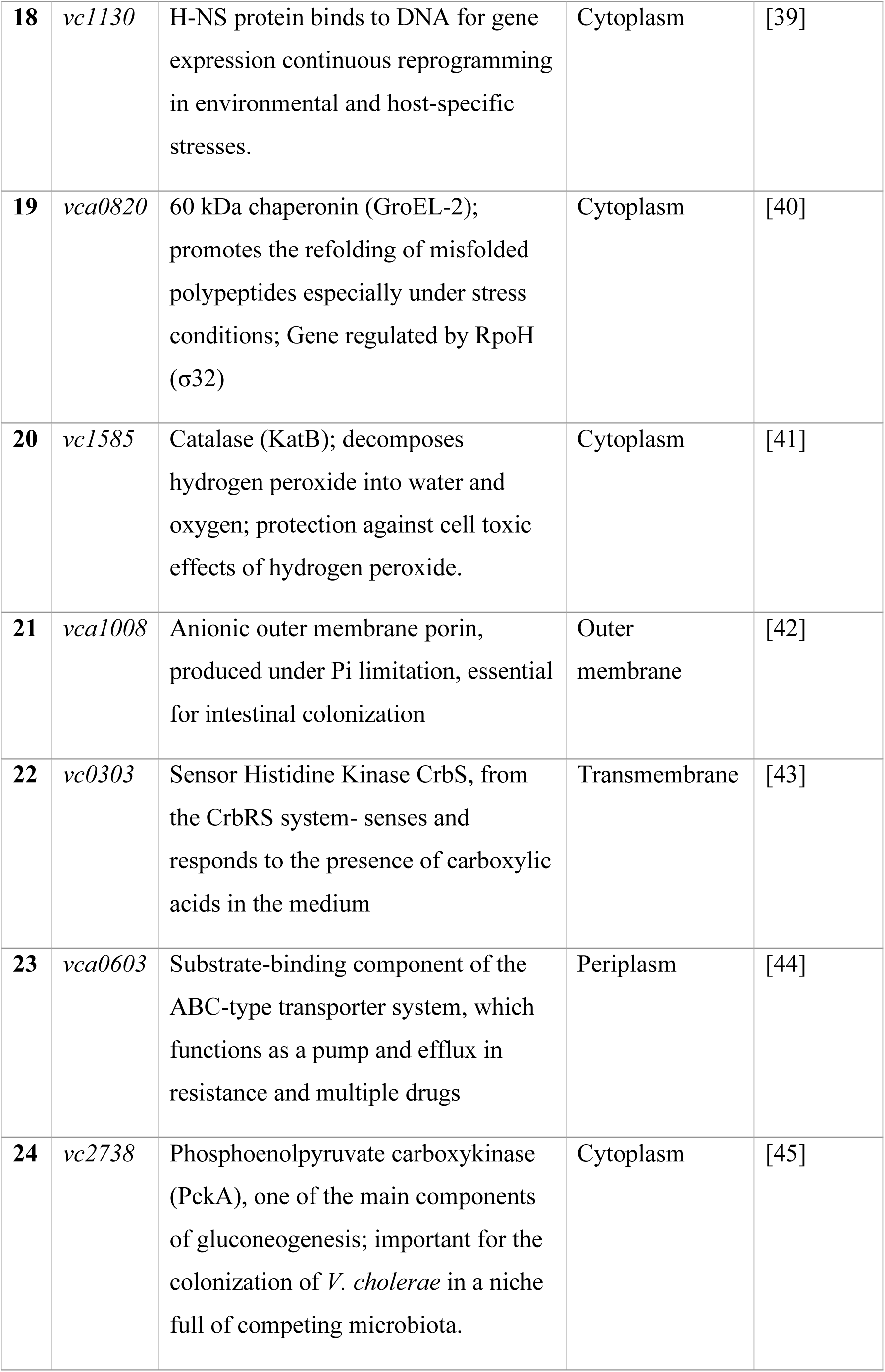
Proteins uniquely identified in OMVs from *Vibrio cholerae* N16961 under phosphate limitation.

The different proteins on Table 1 were mainly derived from the bacterial cytoplasm (75%), followed by 13% from the periplasmic space, 8% from the cytoplasmic membrane, and 4% from the outer membrane. According to their annotated functions, the majority were involved in metabolic pathways (67%). Additionally, 12.5% were related to transport, 12.5% were stress response proteins, and 8% were involved in gene expression regulation.

Genes encoding some of these proteins were found to have PhoB/PhoR-dependent expression, which involves the binding of the response regulator PhoB to conserved 18-bp DNA sequences, called Pho boxes, in the gene regulatory region to activate expression. These data indicate that extracellular Pi limitation not only increases the rate of OMV formation by the bacterium (Fig. 2A) but also regulates the transcription of genes encoding OMV constituent proteins.

Proteins involved in phosphate metabolism, such as the periplasmic Pi-binding protein PstS (VCA0070), alkaline phosphatase PhoA (VCA0033), and the anionic porin PhoE (VCA1008), are essential for nutrient acquisition and intestinal colonization [42,46]. Additionally, enzymes associated with oxidative stress response, including KatB and VC0873, may enhance bacterial survival under Pi-depleted conditions [41]

Beyond phosphate metabolism, *V. cholerae* employs multiple survival strategies and means to establish infection. The synthesis of polyhydroxyalkanoates (PHA), mediated by the enzymes PhaC and PhaB, contributes to carbon storage and stress resistance, as well as playing a role in intestinal colonization [47]. The expression of the *vca0688* gene, which encodes PhaC, is essential for colonization of the mouse intestine, while the PhoB/PhoR system regulates the expression of proteins such as PhaP, which are associated with bacterial responses to phosphate scarcity.

VCA0820 (GroEL) is anothe*r V. cholerae* stress resistance-related protein identified in OMVs released by strain N16961 during growth in low Pi. GroEL is a bacterial chaperonin that, together with its cofactor GroES (GroEL/GroES), plays an essential role in the control of cellular stress, by promoting refolding of misfolded polypeptides and protection to folding protein chains from misfolding and degradation [48].

Proteins such as MalQ, PckA and GltA are essential for carbon metabolism and *V. cholerae* adaptation to changes between aquatic and intestinal environments. These proteins are involved in glycogen synthesis and metabolism, gluconeogenesis, and the citric acid cycle—processes essential for the survival of *V. cholerae* and its successful colonization of the host. In particular, VC2738 (PckA), a phosphoenolpyruvate carboxykinase, and VCA1008 have a critical role in intestinal colonization, being fundamental for *V. cholerae* growth and maintenance in the intestine *in vivo* [42,49].

In summary, our findings on unique proteins packaged in vesicles derived from *V. cholerae* under low Pi suggest a role in phosphate metabolism regulation, oxidative stress response, and host intestinal adaptation mechanisms, all interconnected through a complex gene regulation system mediated by PhoB/PhoR. Importantly, the contribution of these vesicles to pathogenesis is supported by the Galleria mellonella infection assays, demonstrating that factors within OMVs can affect larvae survival. These results indicate a functional link between phosphate metabolism, stress response, and vesicle-mediated effects on the larvae model; however, the precise molecular mechanisms remain to be elucidated.

### Lipidic composition of OMVs liberated by N16961 and Δ*phoB* under limitation and abundance of inorganic phosphate (Pi)

Since bacteria modulate lipid composition to cause bulging leading to OMVs secretion [12], we decided to investigate OMVs lipid composition to correlate it to OMV production.

Outer membrane vesicles released by *Vibrio cholerae* contain a diverse lipid profile, with phosphatidylethanolamine (PE) and ornithine lipid (OL) being the most abundant species. Interestingly, OL, which lacks phosphorus and has two acyl groups linked to ornithine, was found to be the most prevalent lipid in OMVs released by the wild-type strain under low inorganic phosphate (Pi) conditions (Fig. 6). This aligns with expectations, as *V. cholerae* conserves inorganic phosphate in Pi-limited environments by substituting some phospholipids with OL. This process relies on the PhoB/PhoR system, given that *vc0489*, the gene encoding the bifunctional acyltransferase OlsF responsible for OL synthesis, is part of the bacterium’s Pho regulon [50].

**Fig 6.**
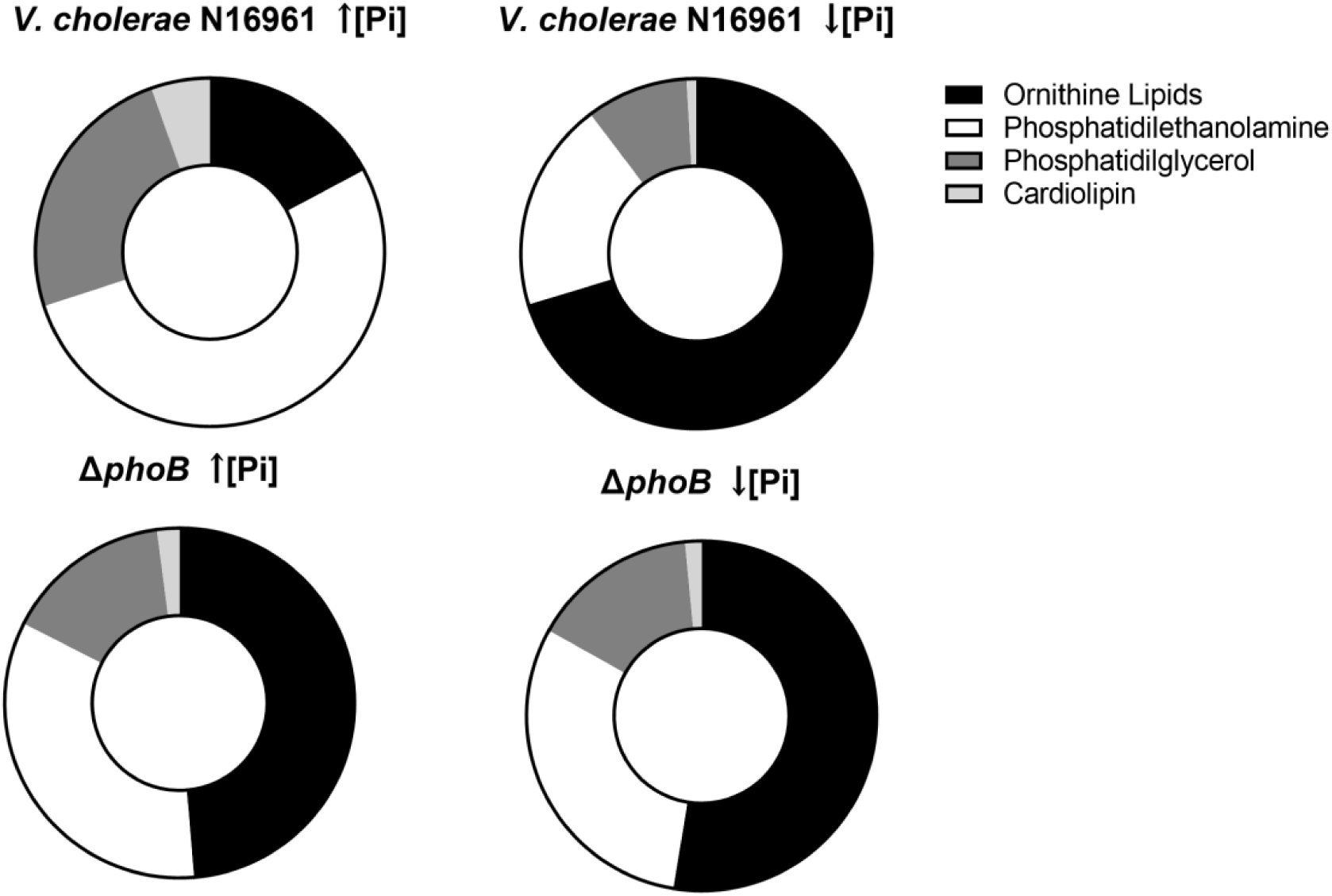
Lipidomic profiles of OMVs from *V. cholerae* N16961 and ΔphoB. Relative distribution of lipid classes in OMVs released by N16961 and ΔphoB under high and low Pi conditions. Lipids were identified by mass spectrometry using LIPID MAPS for phospholipids (PE, PG, CL) and a custom database for phosphate-free ornithine lipids (OL).

Even under high Pi conditions and in the Δ*phoB* mutant (both high and low Pi), OL was detected in OMVs, albeit in lower percentages (<20%). This can be explained by the presence of another gene involved in OL synthesis, *vca0646*, that is expressed in low salinity conditions, which was the case for the media used in these experiments [51].

Bacterial membrane lipid composition plays a crucial role in OMV formation. The distinct physicochemical properties of different lipids influence how they interact to induce outer membrane curvature and facilitate OMV budding (Fig. 1A). While few studies have reported the presence of OL in OMVs, *Bordetella bronchiseptica*, a pig pathogen, is known to release significant amounts of this lipid in its OMVs [11].

However, the interaction of these OL-rich OMVs with the host remains unexplored. *Agrobacterium tumefaciens* also produces phosphorus-free lipids, including OLs, under extracellular phosphate limitation [52]. Lipid analysis of *A. tumefaciens* OMVs revealed that, with the exception of cardiolipin (CL), all other lipids found in bacterial cells were present in the OMVs, though their relative proportions differed between OMVs and cells [53].

Changes in membrane lipids, such as the replacement of zwitterionic PE with positively charged OLs under Pi limitation, can affect interactions with other components like LPS and lipoproteins. This can lead to increased stress and enhanced vesiculation [9].

Consistent with this, both the N16961 wild-type strain and the Δ*phoB* released more OMVs in low Pi conditions compared to high Pi. However, this observation cannot be solely attributed to the quantity of OLs in the original bacterial cell membranes.

In Pi-depleted environments, *V. cholerae* significantly alters its membrane lipid composition to conserve inorganic phosphate for critical metabolic processes (Barbosa et al., 2018). Ornithine lipids are known to suppress the TLR4 response from LPS in immune cells, with their signaling sharing similarities with anti-inflammatory LPS produced by Gram-negative bacteria. This LPS class function is to suppress the host immune system’s TLR4 response, thereby promoting pathogenicity. We propose that the high concentration of ornithine lipids in *V. cholerae* vesicles may contribute to their role in bacterial virulence and host immune system evasion. In phosphate-depleted environments, OLs might replace the immunogenic capacity of bacterial LPS, while constitutively produced OLs could enable bacteria to evade immune surveillance—a known function of OMVs in Gram-negative bacteria [54]

## Conclusion

*Vibrio cholerae* OMV production is sensitive to phosphate availability, being significantly increased under low Pi conditions. Under these conditions, the vesicles not only carry proteins and lipids essential for stress adaptation and nutrient acquisition but also concentrate virulence factors that enhance their toxicity in the *Galleria mellonella* animal model.

The PhoB/PhoR regulon plays a central role in the vesicle cargo loading, determining both the quantity and composition of OMVs. Ornithine lipids, abundant under low Pi, may aid in phosphate conservation and immune evasion, while PhoB/PhoR related proteins can contribute to metabolism, oxidative stress response and pathogenesis.

Our findings suggest that phosphate limitation acts as an environmental signal that selectively increases OMV production and shapes OMV cargo, promoting *V. cholerae* survival, adaptation, and pathogenicity. This study expands the understanding of OMVs as strategic tools for bacterial interaction with the environment, the host and in pathogenicity.

## Methods

### Bacterial strains and OMV Isolation

The culture media used for cell maintenance were LB (Lysogenic broth; 1% bacto-tryptone; 0.5% yeast extract; 1% NaCl) or 1.5% LB-agar (SAMBROOK et al., 1989). For cultivation at known concentrations of inorganic phosphate (Pi), MG (MOPS-glucose) medium was used: MOPS 165 mM pH 7.4, containing a mixture of salts (80 mM NaCl; 20 mM KCl; 20 mM NH_4_Cl; Na_2_SO_4_ 3 mM; 1 mM MgCl_2_; 0.2 mM CaCl_2_), 0.2% glucose, 0.01 M thiamine and KH_2_PO_4_ at final concentrations of 6.5 μM for low-level growth or 65 mM for high-level growth high of Pi [23]. When necessary, streptomycin (Sr, 100 µg/mL) and/or kanamycin (Kan, 50 µg/mL), were added to the media.

N16961 and Δ*phoB* strains were cultivated, respectively, in LB/Sr and LB/Sr.Kan, at 37 °C, under agitation, for 14 h. The next day, a volume of each culture corresponding to OD_600nm_ ∼1.0 was centrifuged and the cell pellet was resuspended in 10 mL of MG. The suspension was diluted 1:100 in 1 L of High Pi/Sr or Low Pi/Sr for the growth of the N16961 strain or in high Pi/Sr.Kan and Low Pi/Sr.Kan, for the cultivation of the Δ*phoB* strain and the flasks were incubated for 14 h at 37 °C under agitation.

The cultures were then centrifuged at 16,000 x g for 10 minutes at 4 °C, the culture supernatants were filtered through a 0.45 µm filter and then through a 0.22 µm filter. The filtrates were subjected to tangential flow filtration on the Vivaflow 200 device (Sigma Aldrich), using a 100 kDa membrane for concentration up to 20 mL. The recovered volume was ultracentrifuged at 120,000 x g for 60 minutes. The supernatants were discarded and the pellet, containing the purified OMVs, was individually resuspended in PBS. Protein content was quantified by the Qubit dosing kit (Thermo Fisher Scientific), following the manufacturer’s instructions. Purified vesicle samples were stored at -20°C.

### Transmission Electron Microscopy

An aliquot (5 µL) of OMV suspension in PBS was applied onto 200 mesh copper grids coated with formvar/carbon film for 1 minute to adsorb the vesicles to the film. As a contrasting agent, 2% uranyl acetate was used for 1 minute and the excess was removed with a paper filter. After air-drying, the samples were observed in the transmission electron microscopes Jeol 1200 EX or Hitachi HT 7800, both operating at 80 kV, at CENABIO-UFRJ.

### Atomic Force Microscopy

An aliquot (5 µL) of OMV suspension was applied onto mica and was dried under an N_2_ (g) atmosphere for 10 min. The samples were analyzed in atomic force microscope Dimension Icon atomic force microscope (Bruker, USA) at CENABIO-UFRJ. The images were acquired in PeakForce QNM® mode at room temperature using a cantilever with a nominal spring constant of 3 N/m, nominal resonance frequency of 75 kHz, and a length of 225 μm. Images were obtained with a resolution of 512 × 512 pixels. The images were processed using the NanoScope Analysis 1.7 software (Bruker).

### Quantification of OMVs by nanoparticle tracking analysis (NTA)

OMV suspension in PBS was used for quantification and characterization using the Nanoparticle Tracking Analysis (NTA) technique under the microscope, with laser and video scattering. For particle tracking the ZetaView® Particle Tracking Analyzer (Particle Metrix, GmbH) or Nanosight NS300 (Malvern Panalytical, Inc.) were used.

The population was defined based on the size, measured by the diameter in nanometers, and the number of vesicles per mL of sample.

### Inoculation of *Galleria mellonella* larvae and assessment of mortality

Larvae of approximately 2 cm in length and 250 mg were used, between the sixth and seventh instar of development, which were individually inoculated in the ventral portion, between the penultimate and last pair of false legs (prolegs) [24,25] with purified OMVs (1 μg of proteins in 10 μL of PBS) derived from cultures in High Pi or Low Pi. Before each inoculation, the larvae were individually transferred to microtubes, which were placed on ice for about 5 min, to anesthetize and facilitate handling and inoculation. The different groups of larvae were separated in glass Petri dishes and incubated in the dark at 30°C. Survival/mortality was recorded every 12 hours for 72 hours. To differentiate between dead and live larvae, the color was initially checked, the dead ones were dark due to the accumulation of myelin. Furthermore, when touched, the dead larvae did not move and when placed in a supine position (with the false paws facing upwards), they were unable to return to the prone position. Ten larvae were inoculated for each type of OMV, and two controls: larvae inoculated with 10 μL of sterile PBS (n = 10 larvae), and another, non-inoculated (n = 10 larvae). 60 larvae were used in total in each independent experiment. To compare the survival rate of larvae from different groups, the Kaplan-Meier method was used, with p values < 0.05 considered statistically significant.

### Extraction of Proteins from OMVs and Identification by LC-MS

A total of 100 µg of OMV proteins were extracted by precipitation with 10% trichloroacetic acid (TCA) then incubated with 0.1% Rapigest SF surfactant (Waters Corporation, USA) and 5 mM dithiothreitol (DTT) to reduce disulfide bridges, for 30 minutes at 60 °C. After cooling, the samples were alkylated with iodoacetamide at a final concentration of 15 mM and then subjected to a dark environment for 30 minutes at room temperature. Each sample was incubated with 1:50 (w/w) of a trypsin solution (Porcine Trypsin, Promega) in 25 mM ammonium bicarbonate pH 8.0, for 18 hours at 37 °C, for protein digestion. The resulting peptide solutions were then dried under vacuum in a SpeedVac for 30 minutes and desalted with ZipTip C18 (Millipore) according to the manufacturer’s instructions. Each sample of desalted tryptic peptides was resuspended in 200 µL of 1% formic acid and 4 µL of this preparation were initially applied to a 2 cm long pre-column (100 µm internal diameter) packed with the 200 A Magic C18 AQ matrix of 5 µm (Michrom Bioresources, Auburn, C.A., USA). The peptides in each sample were then separated into a 10.5 cm long column (75 µm internal diameter) packed with 200 A Magic C18 AQ resin directly over the 15 µm PicoTip (New Objective, Cambridge, MA, USA). For analysis of samples on this column, a flow rate of 2000 nL/min was used and for chromatographic separation, a flow rate of 200 nL/min. Mobile phase A was 0.1% (v/v) formic acid in water, while phase B was 0.1% (v/v) formic acid in acetonitrile, and the gradient conditions were as follows: 2 to 40% B in 57 minutes; up to 80% B in 4 minutes, maintaining this concentration for another 2 minutes, before re-equilibrating the column.

Chromatography was performed using the EASY-nLC II equipment (Thermo Fisher Scientific). The eluted peptides were introduced directly into the nanoLTQ/Orbitrap XL mass spectrometer (Thermo Fisher Scientific) on the Fiocruz/RJ proteomics platform.

The voltage source was 1.9 kV, capillary temperature 200 °C and lens tube voltage was 100 V. The target values for full Ion trap and MSn AGC (Automatic Gain control) were 30,000 and 10,000 respectively, while the FTMS (Fourier Transform Mass Analyzers) AGC target value was set to 500,000. The MS1 spectrum was acquired on the Orbitrap analyzer (m/z 300 to 1700) with 60,000 resolution (m/z 445.1200). For each spectrum, the 10 most intense ions were subjected to CID (collision induced dissociation) fragmentation, followed by acquisition on MS2 of the linear analyzer.

All OMV protein mass spectrometry data were analyzed in triplicates using PEAKS Studio version X software (Bioinformatics Solutions Inc). First, the raw data were submitted to refinement, where the precursor mass and the peak centroiding, charge deconvolution and deisotoping processes were corrected. For further analysis, the tolerance limits were used for the mass values of the precursor ions (in ppm) and for the fragment ions (in Da) (respectively, 10 ppm and 0.5 Da). In addition, enzymatic digestion of the tryptic type was considered and a maximum of two miscleavages per peptide were accepted. All data were initially submitted to the search using the De Novo tool, allowing variable modifications in cysteine (+57.02 Da – carbamidomethylation) and methionine, histidine and tryptophan (+15.99 Da – oxidation), with a maximum of two modifications per peptide. Then, the search was performed using the PEAKS DB search tool, establishing the mass of the precursor as monoisotopic. All searches were performed using the UniProt platform database for *Vibrio cholerae* O1, El Tor biotype, N16961. UniProt Consortium. *Vibrio cholerae serotype O1 (strain ATCC 39315 / El Tor Inaba N16961)*. UniProtKB. Accessed on November 19, 2020. Available at: https://www.uniprot.org/proteomes/UP000000584. The False Discovery Rate (FDR) range was estimated using the decoy tool. Finally, data from triplicates were consolidated and only results in which the estimated FDR was less than or equal to 1% were accepted as reliable identifications.

### Bioinformatic analysis of OMV proteins

Protein sequences identified by mass spectrometry were first annotated using the UniProt database for *Vibrio cholerae* (strain N16961). To predict functional categories, sequences were submitted to the ProtFun 2.2 program, which assigns proteins to Clusters of Orthologous Groups (COGs) based on functional predictions (http://clovr.org/docs/clusters-of-orthologous-groups-cogs/). For subcellular localization, protein sequences were analyzed using the CELLO v2.5 program (http://cello.life.nctu.edu.tw/), which predicts the likely cellular compartment for each protein. Percentages of proteins in each functional category or cellular location were calculated relative to the total number of proteins identified under each experimental condition.

### Extraction of Lipids from OMVs and Identification by LC-MS

Lipids from OMVs were extracted as previously described [55]. In summary, 300 µg of purified and quantified OMVs were resuspended in water (1 mg/mL) and then deuterated standards of PE (phosphatidylethanolamine), PG (phosphatidylglycerol), PC (phosphatidylcholine), and CL (cardiolipin) were added to achieve a final concentration of 0.02 mg/mL. For each mL of the suspension, 3.75 mL of a chloroform and methanol solution (1:2) were added, followed by the addition of 1.25 mL of chloroform and then 1.25 mL of water, with vigorous shaking after each addition. The suspension was centrifuged at 16,000 x g for 20 minutes at 4 °C, the lower phase was transferred to another tube and then dried with nitrogen gas.

Total lipids extracted from the OMVs were then analyzed by mass spectrometry. After extraction, the dried lipids were resuspended in 200 μL of a chloroform and isopropanol mixture (1:4). Ten microliters of each sample were injected in a rapid resolution liquid chromatography (LC) system (1200 series from Agilent Technologies) fitted with a Zorbax XDB Eclipse Plus column (C18, 4.6 × 50 mm, 1.8-μm particle size). The run was 30 min long with the following characteristics: flow rate, 0.3 mL/min; column temperature, 40 °C. Mobile phase A was 0.1% formic acid (positive MS analysis) or 5 mM ammonium acetate, pH 5 (negative MS analysis), and mobile phase B was an isopropanol gradient, which started with 90% solvent A, directly increased to 20% in 10 min, stayed at 20% solvent A for 15 min, and was re-equilibrated to starting conditions in 5 min. A 6520 series electrospray ion source (ESI)-quadrupole time-of- flight (QTOF) high resolution mass spectrometer (Agilent Technologies) was used. For the MS/MS analyses, auto-MS/MS mode was used, and parameters were as follows: positive or negative mode; high resolution acquisition mode (4 GHz); gas temperature, 330 °C; drying gas, 7 liters/min; nebulizer pressure, 50 p.s.i.g.; capillary voltage, −4500 V; fragmentor, 210 V; fixed collision energy, 25 eV; MS scan range and rate, m/z 100–1700 at four spectra per second; MS/MS scan range and rate, m/z 50–1700 at three spectra per second; auto- MS/MS, three maximum precursors; precursor absolute threshold, 200 counts; active exclusion on two repeats and released after 0.5 min. Data were acquired by Mass Hunter Acquisition® for TOF and QTOF version B.04 SP3 (Agilent Technologies). For quantification, single MS analyses were performed with the following parameters: positive or negative mode; extended dynamic range mode (2 GHz); gas temperature, 330 °C; drying gas, 7 liters/min; nebulizer pressure, 50 p.s.i.g.; capillary voltage, 4500 V; fragmentor, 210 V; MS scan range and rate, m/z 100–1700 at two spectra per second. Data was acquired by Mass Hunter Acquisition for TOF and QTOF version B.04 SP3.

The program used for spectrum analysis and database searching was Agilent Mass Hunter (Agilent Technologies). The database used for the general search was obtained from http://www.lipidmaps.org/, and lipids containing ornithine (OLs) were identified using an in-house database created for this purpose.

## Acknowledgements

This work was supported by grants from the Conselho Nacional de Desenvolvimento Científico e Tecnológico (grant number: 408525/2023-1)and the Fundação Carlos Chagas Filho de Amparo à Pesquisa do Estado do Rio de Janeiro. M.L.F. was supported by a fellowship from CAPES. We acknowledge Eduardo Camacho, technician at the Laboratory of Biological Physics, for his valuable technical assistance. We thank the Mass Spectrometry Platform of Fiocruz Rio de Janeiro for analytical support. The authors also acknowledge the National Center for Bioimaging (CENABIO) for providing the infrastructure essential for data acquisition.

## Supporting information

**S1 Table.** Identified proteins in OMVs by LC-MS/MS

